# Cleavage kinetics of human mitochondrial RNase P and contribution of its non-nuclease subunits

**DOI:** 10.1101/2023.03.27.534089

**Authors:** Elisa Vilardo, Ursula Toth, Enxhi Hazisllari, Walter Rossmanith

## Abstract

RNase P is the endonuclease responsible for the 5’-end processing of tRNA precursors (pre-tRNAs). Unlike the single-subunit protein-only RNase P (PRORP) found in plants or protists, human mitochondrial RNase P is a multi-enzyme assembly that in addition to the homologous PRORP subunit comprises a methyltransferase (TRMT10C) and a dehydrogenase (SDR5C1) subunit; these proteins, but not their enzymatic activities, are required for efficient pre-tRNA cleavage. Here we report a detailed kinetic analysis of the cleavage reaction by human PRORP and its interplay with TRMT10C-SDR5C1 using a comprehensive set of mitochondrial pre-tRNAs. Surprisingly, we found that PRORP alone binds pre-tRNAs with nanomolar affinity and can even cleave some of them at reduced efficiency without the other subunits. Thus, the ancient binding mode, involving the tRNA elbow and PRORP’s PPR domain, seems retained by human PRORP, and its metallonuclease domain is in principle correctly folded and functional. Our findings support a model according to which the main function of TRMT10C-SDR5C1 is to direct PRORP’s nuclease domain to the cleavage site, thereby increasing the rate and accuracy of cleavage. Human PRORP’s dependence on the extra tRNA binder appears to have evolved to secure specificity in the cleavage of the structurally degenerating mitochondrial tRNAs.

## INTRODUCTION

In human mitochondria, RNAs are transcribed as long polycistronic precursors, from which mRNAs, rRNAs, and tRNAs have to be released by endonucleases before further maturation to their final functional forms. Due to the interspersed and butt-jointed arrangement of the tRNA sequences, the two enzymes RNase P and RNase Z, responsible for 5’- and 3’-end processing of tRNAs respectively, concomitantly generate the 3’ and 5’ ends of the two rRNAs and most mRNAs, and thereby account for almost the entire primary processing of mitochondrial transcripts (1–4). While mitochondrial RNase Z is derived from the same gene (*ELAC2*) as its nuclear isoform (1,5,6), mitochondrial RNase P (mtRNase P) is different from and unrelated to nuclear RNase P (7–9). Remarkably, mtRNase P is also a multi-enzyme assembly that appears exceptional, even within the diverse RNase P enzyme family (10). In the eukaryal domain the RNase P family encompasses ribonucleoproteins of various levels of complexity and unrelated composition, as well as a class of comparatively “simple”, monomeric 60-kDa protein enzymes called <u>pro</u>teinaceous <u>R</u>Nase <u>P</u> (PRORP) (10,11). PRORP proteins function as RNase P in the nucleus, mitochondria, and/or plastids of land plants, algae, trypanosomes, and probably several other eukaryal groups (12–18). Also human mtRNase P comprises a PRORP protein (aka MRPP3) as endonuclease, but it requires two additional essential protein subunits, which have further, unrelated functions (9,19): short-chain dehydrogenase/reductase family 5C member 1 (SDR5C1 aka MRPP2 or HSD17B10) catalyzes the penultimate step in the β-oxidation of short branched-chain fatty and amino acids (20), and tRNA methyltransferase 10C (TRMT10C aka MRPP1) forms a stable subcomplex with SDR5C1 that constitutes the methyltransferase responsible for *N*^1^-methylation of purines at position 9 of mitochondrial tRNAs (19).

We previously showed that SDR5C1 is not directly involved in tRNA binding and neither its dehydrogenase activity nor an intact binding site for its NAD^+^/NADH cofactor are required for the methyltransferase or endonucleolytic activity of the mtRNase P complex, suggesting a scaffolding role for SDR5C1 (10,19,21). Similarly, the methyltransferase activity of TRMT10C is dispensable for cleavage by PRORP, supporting a tRNA-binding and structural, rather than enzymatic role of the TRMT10C-SDR5C1 complex in promoting the cleavage reaction (19). Recently, the cryo-electron microscopy (cryo-EM) structure of human mtRNase P bound to precursor tRNA (pre-tRNA) was reported, showing that TRMT10C encases part of the tRNA, and forms several specific and nonspecific interactions with all four subdomains of the tRNA structure (22). TRMT10C is anchored to the SDR5C1 tetramer via a central loop and helix connecting its two tRNA-interacting domains, while SDR5C1 itself appears to have limited contact with the anticodon loop of the tRNA. An arch-like shaped PRORP binds on top of the pre-tRNA-TRMT10C-SDR5C1 complex through interactions with both, the pre-tRNA and TRMT10C; its nuclease domain contacts the pre-tRNA around the cleavage site and the methyltransferase domain of TRMT10C, and on the other end, its pentatricopeptide repeat (PPR) domain interacts with the elbow of the tRNA structure and the N-terminal domain of TRMT10C. Active-site organization and the overall geometry of the PRORP-tRNA interaction appear similar to that of (single-subunit) PRORPs from *Arabidopsis thaliana* or of their models in complex with tRNA (23–26), yet the molecular interactions between the PPR domain and the tRNA elbow appear to be different (22,27).

The structure of the mtRNase P-pre-tRNA complex confirmed the originally proposed role of the TRMT10C-SDR5C1 complex as an accessory factor in the recognition of pre-tRNAs (9,10,22). However, a suggested role in remolding the disordered catalytic domain observed in the crystal structures of PRORP fragments, to enable the coordination of Mg^2+^ in the latter’s active site, appears not entirely consistent with the reported structural and enzymatic data (22,28,29). Thus, the actual mechanisms behind the interplay between TRMT10C-SDR5C1 and PRORP leading to pre-tRNA cleavage remain unclear.

Here we report a detailed analysis of tRNA 5’-end maturation by human mtRNase P using a comprehensive and representative set of mitochondrial pre-tRNA substrates. We investigated the contribution of the individual mtRNase P subunits to 5’-end processing, their pre-tRNA-binding kinetics, and the cleavage process itself under a variety of kinetic regimens. We also examined previously raised hypotheses about the role(s) of TRMT10C-SDR5C1. Altogether our results reveal new insights into the mechanism and conformational dynamics of an exceptionally multifaceted member of the diverse RNase P enzyme family.

## MATERIALS AND METHODS

### Expression and purification of recombinant proteins

C-terminally His-tagged human PRORP, C-terminally myc-His-tagged TRMT10C, N-terminally His-tagged SDR5C1, N-terminally His-tagged *Saccharomyces cerevisiae* Trm10p, and C-terminally His-tagged *Arabidopsis thaliana* PRORP3 were prepared as previously described (19,30). The TRMT10C-SDR5C1 complex was essentially prepared as previously described (19), with the imidazole concentration during the wash step increased to 200 nM.

Substitutions D479N and D499N were introduced into the PRORP expression plasmid by site-directed mutagenesis using the QuikChange protocol (Agilent Technologies). The two variants were expressed and purified like the wild type protein (19).

ELAC2, starting with amino acid 16, was cloned the *Nde*I/*Xho*I sites of pET-21b(+) and, including the stop codon, into the same sites of pET-28b(+) (Novagen); thereby the recombinant proteins include either a C-terminal or an N-terminal 6×His-tag. C- and N-terminally His-tagged ELAC2 (indicated as ELAC2-His and His-ELAC2, respectively) were expressed in *Escherichia coli* Rosetta2(DE3) and purified on HisTrap HP columns (Cytiva) essentially as previously described for SDR5C1 (19): proteins were loaded and washed with 50 mM imidazole, then washed with 1 M NaCl, followed by 300 mM imidazole, and eluted with 500 mM imidazole (all in buffer A).

The purity of the recombinant proteins and the stoichiometry of the TRMT10C-SDR5C1 complex were assessed by SDS-PAGE and Coomassie brilliant blue staining, and found to be more than 95% and consistent with a 2:4 ratio, respectively. All protein concentrations were quantitated relative to bovine serum albumin standards by SDS-PAGE, Coomassie brilliant blue staining, and image analysis using ImageQuant TL 8 (Cytiva). In the case of the purified TRMT10C-SDR5C1 complex, the indicated concentrations refer to the concentration of TRMT10C monomers as the presumable active tRNA-binding unit.

### Precursor tRNA substrates

The pre-tRNAs used for RNase P activity assays were human mitochondrial tRNAs with short stretches of their natural 5’- and 3’-flanking sequences. The plasmid constructs for the *in vitro* transcription of pre-tRNA^Tyr^, pre-tRNA^Ile^ and pre-tRNA^His^ were previously described (9,19). Similarly, plasmid constructs for the *in vitro* transcription of further pre-tRNAs were generated by PCR cloning of the following regions of the human mitochondrial genome: 5691–5567 for pre-tRNA^Ala^, 5891–5738 for pre-tRNA^Cys^, 4434–4312 for pre-tRNA^Gln^, 14836–14639 for pre-tRNA^Glu^, 8254–8375 for pre-tRNA^Lys^, 4368–4503 for pre-tRNA^Met^, 16060–15926 for pre-tRNA^Pro^, 7521–7441 for pre-tRNA^Ser(UCN)^, and 1571–1702 for pre-tRNA^Val^. Their *in vitro* transcripts start in most cases with a short stretch of polylinker sequence preceding the natural 5’-flanking sequence of the respective tRNA. The template for the RNase P model substrate *Thermus thermophilus* pre-tRNA^Gly^ was previously described (30,31).

Pre-tRNAs used for RNase Z activity assays were 5’-mature human mitochondrial tRNAs with a short stretch of 3’-flanking sequence only. For pre-tRNA^His^ (mtDNA 12138– 12223), pre-tRNA^Leu(UUR)^ (3230–3340), and pre-tRNA^Ser(UCN)^ (7514–7426), transcription templates were derived by *in vitro* mutagenesis of plasmids encoding tRNAs with 5’- and 3’-flanking sequences; thereby all sequences between the T7/T3 polymerase promoter initiation site and the 5’ end of the tRNA were deleted to start *in vitro* transcription directly with the G1 of the tRNA. In the templates for pre-tRNA^Ala^ (5655–5567) and pre-tRNA^Ile^ (4263–4351), the tRNAs’ 5’ end was fused to a hammerhead ribozyme to release the 5’ end of the tRNA after *in vitro* transcription. The plasmid construct for pre-tRNA^Ala^ was assembled by overlap extension PCR and cloned as previously described (19), the one for pre-tRNA^Ile^ was a kind gift from Louis Levinger (32).

*In vitro* transcription, ^32^P-5’-end labeling and purification of pre-tRNAs were carried out as previously described (7,33). The concentration of unlabeled pre-tRNAs was determined by UV spectrophotometry, the one of labeled pre-tRNAs with the RiboGreen RNA quantitation reagent (Molecular Probes) using standard curves generated by serial dilution of unlabeled pre-tRNA.

### RNase P and RNase Z cleavage assays

Qualitative RNase P cleavage assays were performed at 21 °C as previously described (21). Quantitative kinetic analyses were performed at 30 °C in the previously described mtRNase P reaction buffer (21), but with either 3 or 4.5 mM Mg^2+^, as found optimal for the cleavage of the respective pre-tRNA and indicated in the figure legends. The experimental procedure was detailed previously (30) and the differences are reported in the following.

In single-turnover kinetic experiments (final substrate concentration 0.2 nM), labeled pre-tRNAs and PRORP, each at twice the final concentration, were pre-incubated separately in reaction buffer at 30 °C, and reactions started by combining equal volumes of the two. To analyse PRORP cleavage in the presence of the TRMT10C-SDR5C1 complex (mtRNase P holoenzyme), the latter was added to the substrate at the preincubation step to 400 nM (i.e., 200 nM final concentration, which was verified to be saturating for all tested substrates in pilot experiments). A similar regimen was followed in multiple-turnover kinetic experiments, but the pre-tRNA substrates were pre-incubated with a fourfold molar excess of TRMT10C-SDR5C1 (or at least 200 nM final concentration in the case of the lower substrate concentrations); the different concentrations of unlabeled substrate in multiple-turnover experiments were spiked with labeled substrate at 1 or 2 nM. The concentration of PRORP in multiple-turnover kinetic experiments was 1 nM. Sampling and all further processing of reaction time points were carried out as previously described (30).

Aliquots of single-turnover reactions with the mtRNase P holoenzyme (PRORP plus TRMT10C-SDR5C1) were sampled throughout the reaction and pseudo first-order rate constants of cleavage (*k*_obs_) were calculated by nonlinear regression analysis fitting the data to the equation for a single exponential: *f*_cleaved_ = *f*_endpoint_ × (1 − e^−(*k*obs) × *t*^), where *f*_cleaved_ = fraction of substrate cleaved, *t* = time, *f*_endpoint_ = maximum cleavable substrate (Prism, GraphPad Software). Single-turnover reactions with PRORP alone were considerably slower and, particularly with lower enzyme concentrations, did not reach meaningful endpoints within reasonable incubation times. Therefore, aliquots were sampled throughout the initial, largely linear phase of the reactions only, and linear regression analysis was used to approximate the pseudo first-order rate constants (*k*_obs*_) of cleavage. Similarly, aliquots of multiple-turnover reactions were sampled throughout the initial largely linear phase of the reactions, and linear regression analysis was used to approximate the initial velocity (*V*_0_) of cleavage.

For the analysis of single-turnover kinetics, *k*_obs_ or *k*_obs*_ values from at least six different enzyme concentrations, from at least four replicate experiments each, were plotted against the enzyme concentration; the maximal rate constant (*k*_react_) and the enzyme concentration at which the half-maximal rate constant is achieved (*K*_M(sto)_) were calculated by nonlinear regression, fitting the data to a “Michaelis-Menten-like” enzyme kinetics model: *k*_obs_ = *k*_react_ × [PRORP] / (*K*_M(sto)_ + [PRORP]). For the analysis of multiple-turnover kinetics, *V*_0_ values from at least seven different substrate concentrations, from at least four replicate experiments each, were plotted against the substrate concentration; the turnover number (*k*_cat_) and the Michaelis-Menten constant (*K*_M_) were calculated by nonlinear regression, fitting the data to a Michaelis-Menten enzyme kinetics model: *V*_0_ = [PRORP] × *k*_cat_ × [pre-tRNA] / (*K*_M_ + [pre-tRNA]). The results are reported as best-fit values ± curve-fit standard error (Prism, GraphPad Software).

Pulse-chase reactions were essentially carried out as previously outlined (34). In brief, single-turnover kinetic reactions with 50 nM PRORP and 200 nM TRMT10C-SDR5C1 were set up as described above. After the indicated time the reactions were chased either by the addition of unlabeled pre-tRNA to 1 µM or by 200-fold dilution with reaction buffer. Samples from the diluted reactions were precipitated with ethanol/ammonium acetate/glycogen before dissolving them in the gel loading buffer. Further processing and analysis were carried out as described above.

To determine the optimal Mg^2+^ concentration for pre-tRNA cleavage, enzyme kinetics were carried out at Mg^2+^ concentrations ranging from 0.5 or 1.5 up to 9/12 mM, with 200 nM PRORP alone, or with 20 nM PRORP plus 200 nM TRMT10C-SDR5C1 complex; *k*_obs_ values were derived from at least three replicate experiments.

RNase Z cleavage and kinetics were performed at 30 °C in a buffer composed like the mtRNase P reaction buffer, but with pH 7.4 (because ELAC2 was unstable at pH 8) and 1 mM Mg^2+^. Assays and analyses of single-turnover kinetics were carried out as described above for the mtRNase P holoenzyme.

### RNA binding studies

The binding of the subunits of mtRNase P to different mitochondrial pre-tRNAs was measured by bio-layer interferometry on a BLItz instrument (Pall FortéBio). For immobilization on streptavidin-coated sensors pre-tRNAs were 5’- or 3’-end biotinylated. Briefly, *in vitro* transcribed pre-tRNA^Ala^ and pre-tRNA^His^ (see previous section) were ligated to a 5’-biotinylated DNA oligonucleotide (bio-TGTGGTTGACGTGC) each using a complementary bridging oligonucleotide (splint) and T4 DNA ligase (35). For pre-tRNA^Lys^ and pre-tRNA^Val^, the regions corresponding to nucleotides 9–73 plus 3’-flanking sequence (8303–8375 and 1610–1702 of the mitochondrial genome, respectively) were PCR amplified, assembled by overlap extension with a hammerhead ribozyme at the 5’ end, and cloned into the plasmid pGEM-1 (Promega) as previously described (19). The *in vitro* transcribed tRNA fragments were completed by “splint ligation” to a synthetic RNA corresponding to the first 8 nucleotides of the respective tRNA plus 15 nucleotides of the natural 5’-flanking sequence (19). The pre-tRNAs were 3’ biotinylated with pCp-biotin (750 µM; Jena Bioscience) and T4 RNA ligase (2 units/µl), at 16 °C over night in T4 RNA ligation buffer (100 mM Tris-Cl pH 7.8, 100 mM MgCl_2_, 100 mM DTT, 10 mM ATP). Gel-purified pre-tRNAs were immobilized on streptavidin-coated sensors, and protein binding was assayed upon incubation with at least three different concentrations in the range of 50–1000 nM at room temperature set to 21 °C (the instrument itself does not allow temperature control). All steps were performed in the following binding buffer: 50 mM Tris-Cl pH 8, 20 mM NaCl, 2 mM DTT, 20 µg/ml BSA, 0.5 units/µl Ribolock RNase inhibitor (Fermentas), 0.05% Tween-20 (to minimize unspecific binding to the sensor), with or without 4.5 mM MgCl_2_ (Mg^2+^ was included in the analysis of tRNA^Lys^, but excluded in the case of the other pre-tRNAs to prevent cleavage by PRORP; in pilot experiments, moreover Mg^2+^ did not significantly affect binding by TRMT10C or PRORP). Association and dissociation were recorded for 3–5 and 5–30 minutes, respectively (depending on the respective pre-tRNA-protein interaction studied). The binding data were analysed with the BLItz Pro software (Pall FortéBio) applying step correction and global fitting; results are reported as best-fit values ± curve-fit standard error.

### Iron-mediated cleavage of PRORP

For Fe(II)-mediated hydroxyl radical cleavage (36) 7.2 µM PRORP was incubated with 200 μM (NH_4_)_2_Fe(SO_4_)_2_, 10 mM DTT, and 10 mM ascorbic acid, in 50 mM PIPES-K pH 6.7 buffer. TRMT10C-SDR5C1 complex and/or pre-tRNA^Ile^ (see section above) were added to a final concentration of 7.8 and 7.5 µM, respectively. Reactions were incubated at 20°C for 1 hour, stopped by addition of 5 × Laemmli buffer, and analysed by 12–20%-gradient SDS-PAGE and Coomassie brilliant blue staining. The identity of the C-terminal fragments produced by Fe(II)-mediated cleavage was confirmed by western blotting using an anti-His-tag antibody.

## RESULTS

### PRORP alone is able to cleave pre-tRNAs *in vitro*

With the identification of the subunits of human mtRNase P, we originally found all three proteins to be required for the reconstitution of the pre-tRNA-cleavage activity (9). However, various PRORP homologues later identified in plants or protists were found to have RNase P activity on their own, without the involvement of additional proteins (12–17). The architecture of human PRORP moreover shows all the defining features generally found in PRORP homologues (18), i.e., a characteristic C-terminal NYN metallonuclease domain, an N-terminal α-super helical domain containing pentatricopeptide repeat (PPR) motifs, and a bipartite zinc-binding module connecting these two domains, in a largely identical three-dimensional arrangement (22). We thus decided to re-examine the activity of human PRORP alone in comparison to that of the mtRNase P holoenzyme more thoroughly and on a wider set of human mitochondrial pre-tRNA substrates. Confirming our previous observation (9), pre-tRNA^Ile^ and pre-tRNA^Tyr^, as well as several other mitochondrial pre-tRNAs were not specifically cleaved by PRORP alone even at an enzyme concentration 25-fold higher than the one originally used (Figure 1A). However, in the case of 5 out of the 10 newly-tested mitochondrial pre-tRNAs, we detected substantial RNase P activity by PRORP alone, particularly at elevated concentrations (Figure 1B). Nevertheless, in all those cases the addition of TRMT10C and SDR5C1 caused a strong stimulation of the specific pre-tRNA cleavage. The latter observation is consistent with the strict requirement of all three proteins for efficient 5’-end processing of mitochondrial tRNAs *in vivo*, as previously demonstrated by the accumulation of mitochondrial pre-tRNAs upon knock-down of either TRMT10C, SDR5C1, or PRORP (9). Taken together, these findings indicate that human PRORP harbours a complete, functional active site and is able to recognize, bind, and cleave at least certain pre-tRNAs at the correct site on its own *in vitro*, whereas TRMT10C-SDR5C1 appear primarily required to increase the cleavage efficiency of PRORP.

**Figure 1.**
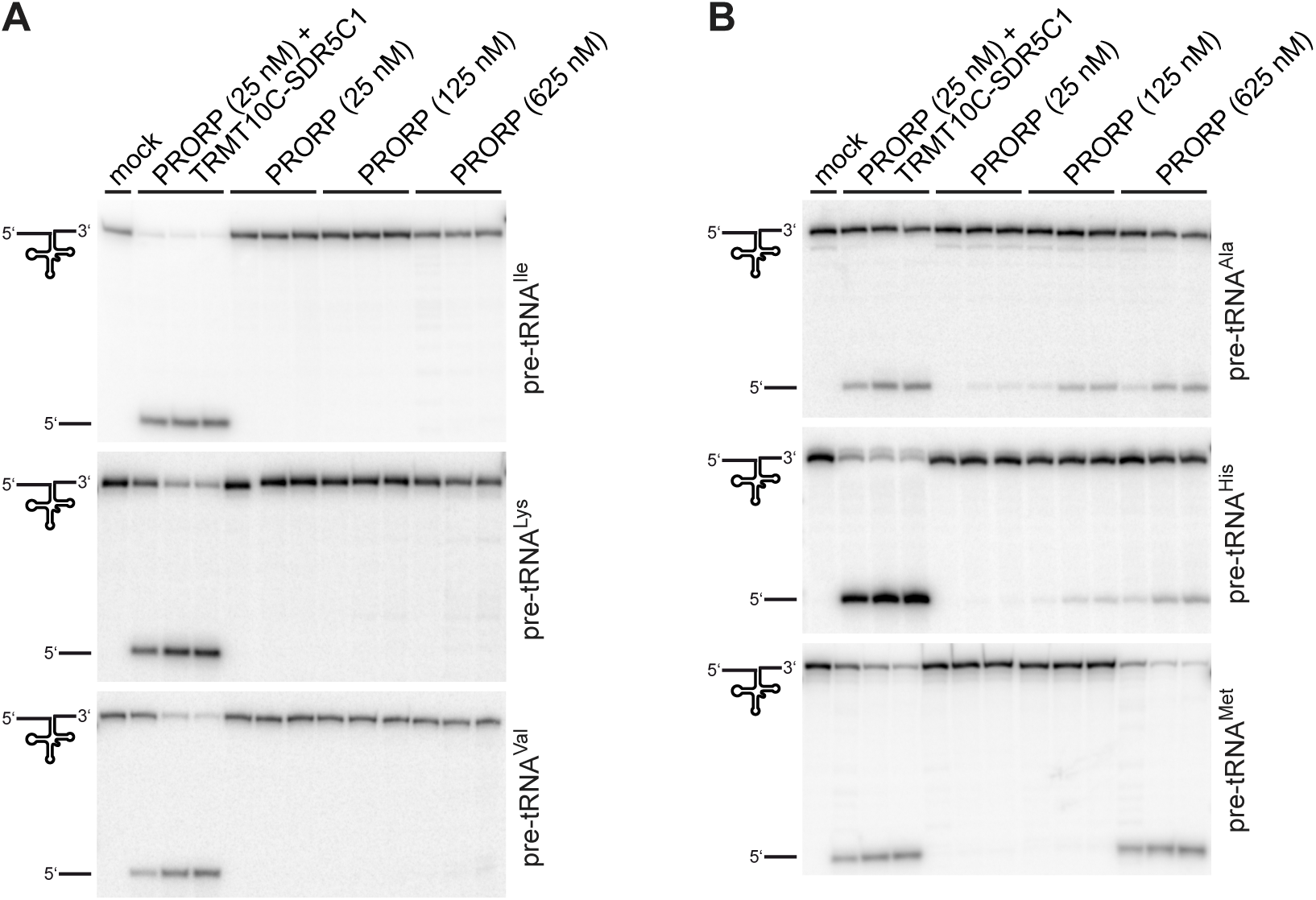
PRORP has RNase P activity without TRMT10C-SDR5C1. The RNase P activity of human PRORP was tested on model substrates of the six indicated human mitochondrial pre-tRNAs. Aliquots were withdrawn from the reactions after 3, 30, and 60 minutes for pre-tRNA^Ala^ and pre-tRNA^Val^, and 10, 30, and 60 minutes for pre-tRNA^His^, pre-tRNA^Ile^, pre-tRNA^Lys^ and pre-tRNA^Met^; cleavage products were separated by gel electrophoresis and visualized by phosphorimaging. The final concentration of the TRMT10C-SDR5C1 complex, when added to the reaction, was 250 nM. No enzyme was added to the “mock” reaction, which was incubated for 60 minutes. Due to the 5’-end labeling, only the full length pre-tRNA and the released 5’ leader are visible. (**A**) Examples of human mitochondrial pre-tRNAs that required both, PRORP and the TRMT10C-SDR5C1 complex for 5’-end processing. (**B**) Examples of human mitochondrial pre-tRNAs on which PRORP alone showed RNase P activity *in vitro*.

Animal mitochondrial tRNAs are structurally heterogeneous, deviate from canonical tRNA structures in the size of their D and TΨC loops, are characterized by a low number of G-C base pairs in the stems, and typically lack one or more of the common tertiary-interaction motifs (37). In addition to 12 mitochondrial pre-tRNAs, we also tested the processing of a fully canonical pre-tRNA by PRORP in the presence or absence of TRMT10C-SDR5C1. We chose pre-tRNA^Gly^ from *T. thermophilus*, a well-characterised bacterial model substrate that was previously used in a large number of studies with a wide variety of RNase P enzymes, including various PRORP homologues (15,30,38–45). The mtRNase P holoenzyme cleaved the bacterial pre-tRNA^Gly^ at the canonical cleavage site between position +1 and −1, like *A. thaliana* PRORP3 and any other previously tested form of RNase P (Figure 2). However, human PRORP alone not only showed weaker cleavage, but about 50% of the cleavage occurred one nucleotide upstream, between −1 and −2, in addition to the canonical site (Figure 2). These results suggest that TRMT10C-SDR5C1 may also affect the correct positioning of the target phosphodiester bond in the active site of PRORP, and that without the aid of TRMT10C-SDR5C1, PRORP’s ability to correctly select the cleavage site on some substrates may be impaired.

**Figure 2.**
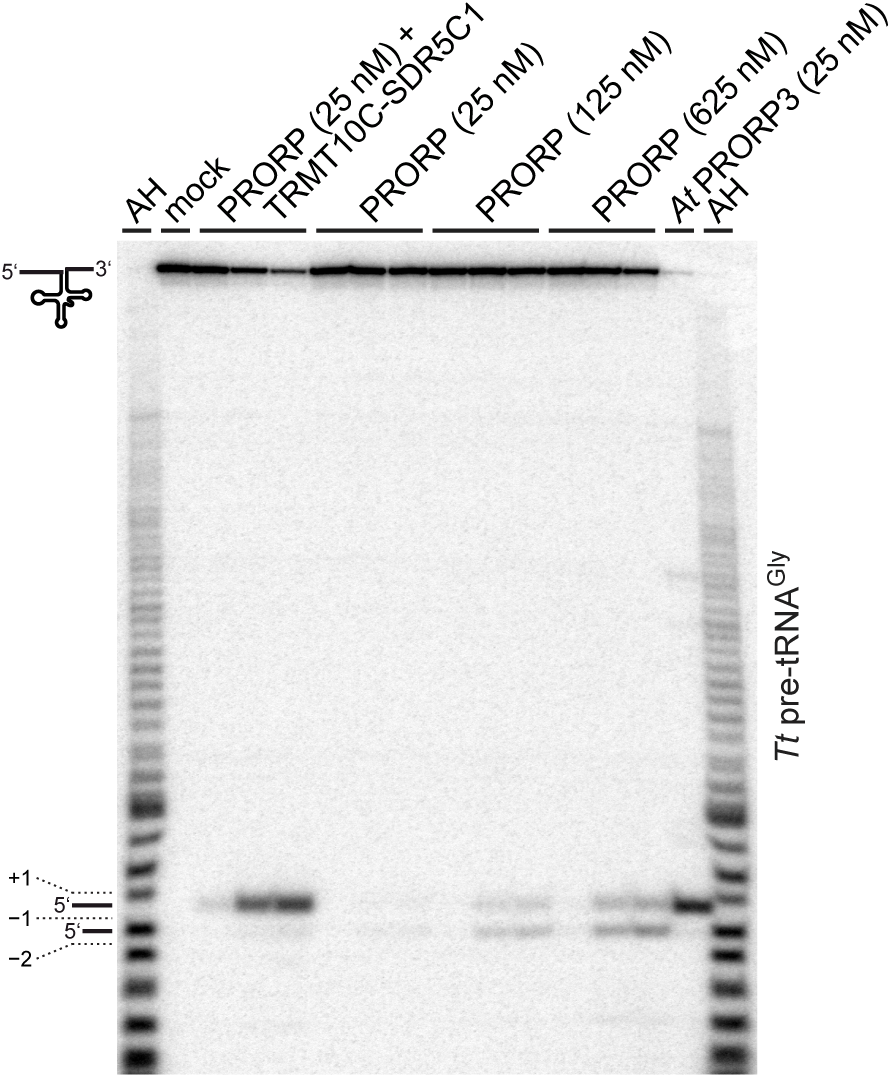
(Mis-)cleavage of an RNase P-reference substrate by human PRORP. The RNase P activity of human PRORP was tested on the bacterial *T. thermophilus* (*Tt*) pre-tRNA^Gly^. Aliquots were withdrawn from the reactions after 3, 30, and 60 minutes, and analyzed as described for Figure 1. In the first and last lanes, an alkaline hydrolysis ladder (AH) generated from pre-tRNA^Gly^ was loaded as migration marker (the off-set in migration is due to the 2’-3’-cyclic phosphates produced by alkaline hydrolysis as opposed to the 3’-hydroxyl ends produced by RNase P). A processing reaction with *A. thaliana* PRORP3 (*At* PRORP3) was included to identify the position of the canonical cleavage site between +1 and −1. (Mis-)cleavage between pre-tRNA nucleotides −1 and −2 results in a one-nucleotide shorter 5’-leader product.

### The effect of TRMT10C-SDR5C1 on pre-tRNA processing is specific to PRORP

The TRMT10C-SDR5C1 complex was previously suggested to act as a general tRNA-maturation platform in mitochondria, a concept based on the observation that besides being a mtRNase P subunit it also appeared to stimulate the cleavage by mitochondrial RNase Z (ELAC2) (46). While postulating a general role of TRMT10C-SDR5C1 in facilitating mitochondrial tRNA processing appears attractive, it is on the other hand difficult to reconcile with the direct interactions of PRORP and TRMT10C as observed in the recently determined mtRNase P-pre-tRNA structure (22). A common underlying mechanism would either require specific protein-protein interactions of largely the same TRMT10C surfaces with both, PRORP and ELAC2, or a more unspecific, probably tRNA structure-mediated effect of TRMT10C-SDR5C1 to similarly stimulate the cleavage by the different effector nucleases. We thus re-assessed the stimulatory effect of TRMT10C-SDR5C1 on the RNase Z activity of ELAC2, and explored the possibility that remolding of tRNA structures by TRMT10C-SDR5C1 (or a related methyltransferase) per se may stimulate cleavage.

In contrast to previous observations (46), TRMT10C-SDR5C1 inhibited, rather than stimulated, the RNase Z activity of ELAC2 in a dose dependent manner (Figure 3A). This result was corroborated for 4 more mitochondrial pre-tRNAs, under multiple as well as single turnover conditions, under higher ionic strength (as used in the previous study), and with recombinant ELAC2 tagged at either the C- or the N-terminus. Noteworthy, the efficiency of pre-tRNA^Ala^ cleavage by ELAC2 alone resembled much more that of the mtRNase P holoenzyme than that of PRORP alone (compare next section, Figure 4).

**Figure 3.**
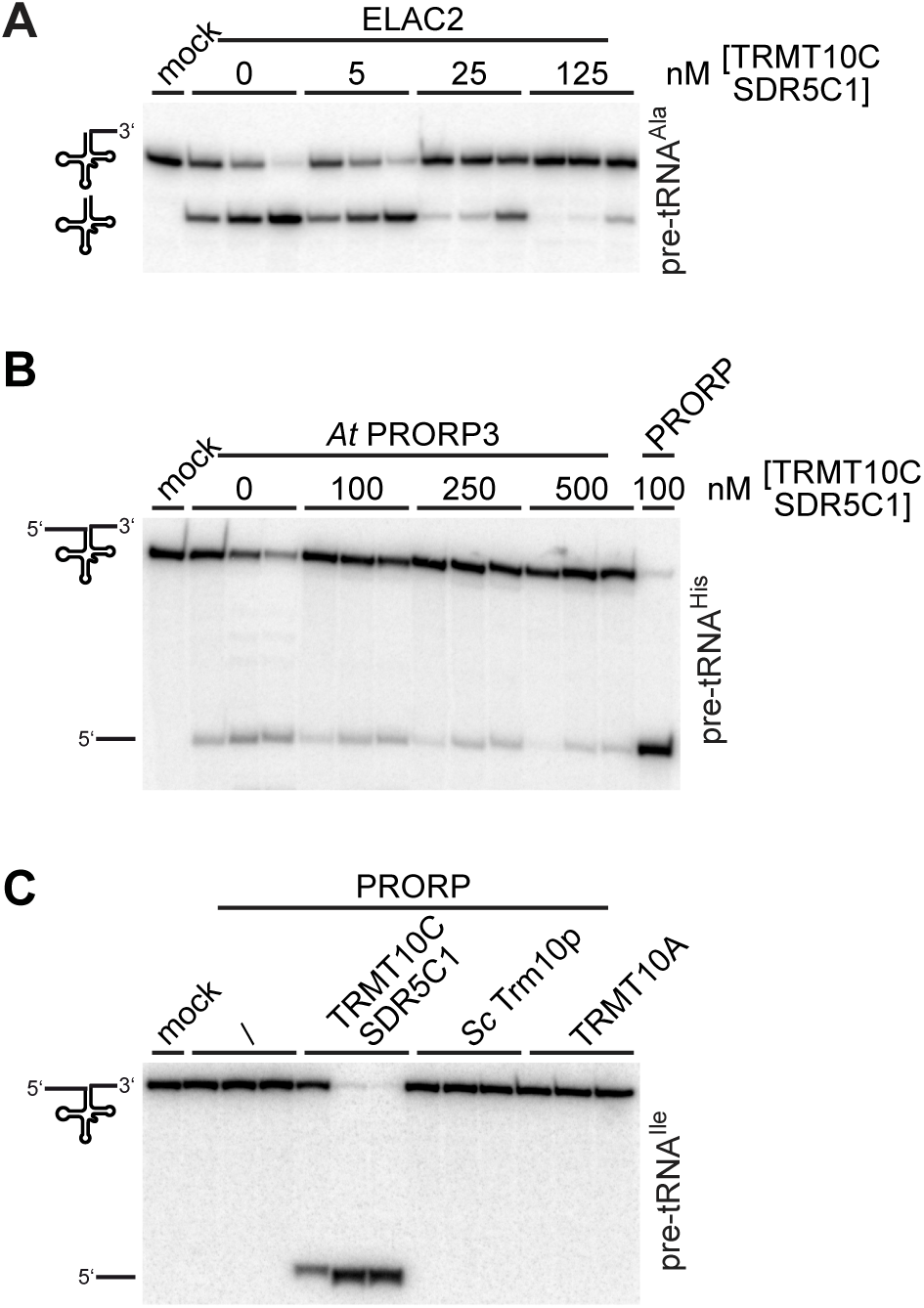
TRMT10C-SDR5C1 inhibits the activity of ELAC2 and *A. thaliana* PRORP3, and cannot be replaced by other TRM10 methyltransferases in mtRNase P. **(A)** The effect of the TRMT10C-SDR5C1 complex on the RNase Z activity of ELAC2 was tested. Mitochondrial pre-tRNA^Ala^ was supplemented with the indicated concentrations of TRMT10C-SDR5C1, and tested for cleavage by His-ELAC2 (25 nM); aliquots were withdrawn at 1, 5, and 60 minutes, and analyzed as described for Figure 1. **(B)** The ability of the TRMT10C-SDR5C1 complex to stimulate RNase P cleavage was tested with a plant PRORP homologue. Human mitochondrial pre-tRNA^His^ was supplemented with the indicated concentrations of human TRMT10C-SDR5C1, and tested for cleavage by *A. thaliana* PRORP3 (*At* PRORP3; 100 nM); aliquots were withdrawn at 3, 30, and 60 minutes, and analysed as described for Figure 1. (C) The RNase P activity of human PRORP (25 nM) was tested on human mitochondrial pre-tRNA^Ile^ in the presence of different methyltransferases of the TRM10 family: the human mitochondrial TRMT10C-SDR5C1 complex, *S. cerevisiae* Trm10p (*Sc* Trm10p), and human nuclear TRMT10A, each at a final concentration of 250 nM. Aliquots were withdrawn after 15 seconds, 15 and 60 minutes, and analyzed as described for Figure 1.

**Figure 4.**
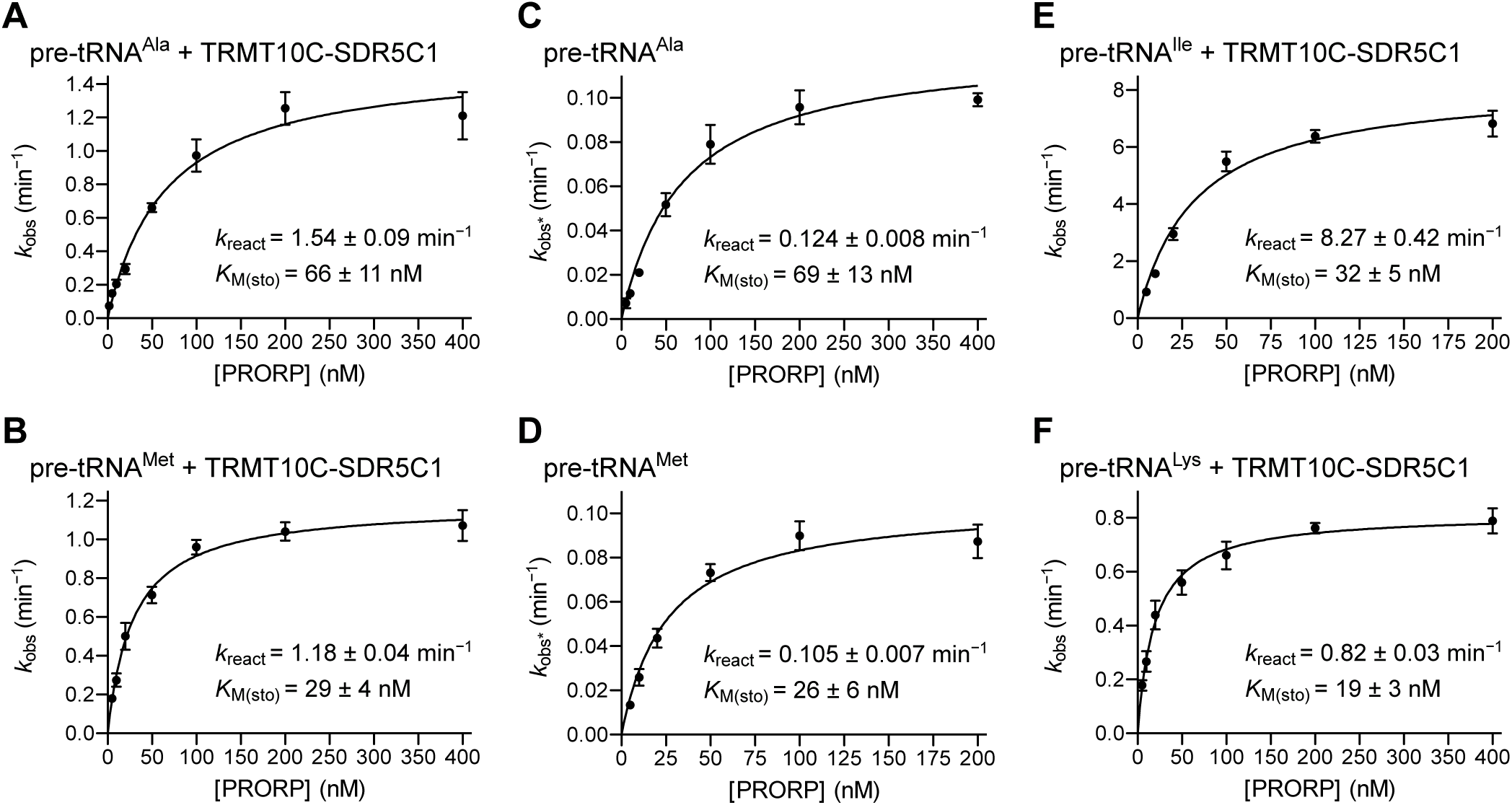
Single-turnover kinetics of pre-tRNA cleavage by the mtRNase P holoenzyme and PRORP alone. Single-turnover kinetic analyses of pre-tRNA cleavage were performed with varying concentrations of PRORP in the presence of an excess of TRMT10C-SDR5C1 complex (mtRNase P holoenzyme; panels A, B, E, and F) or PRORP alone (panels C and D). Pseudo first-order rate constants of cleavage (*k*obs or *k*obs*) were plotted against the concentration of PRORP. Data points are the mean ± SEM of at least 5 replicates. Derived kinetic constants *k*react and *K*M(sto) are inserted into each graph. (**A**) Kinetic analysis of the cleavage of pre-tRNA^Ala^ by PRORP in presence of TRMT10C-SDR5C1 (3 mM Mg^2+^). (**B**) Kinetic analysis of the cleavage of pre-tRNA^Met^ by PRORP in presence of TRMT10C-SDR5C1 (4.5 mM Mg^2+^). (**C**) Kinetic analysis of the cleavage of pre-tRNA^Ala^ by PRORP alone (3 mM Mg^2+^). (**D**) Kinetic analysis of the cleavage of pre-tRNA^Met^ by PRORP alone (4.5 mM Mg^2+^). (**E**) Kinetic analysis of the cleavage of pre-tRNA^Ile^ by PRORP in presence of TRMT10C-SDR5C1 (3 mM Mg^2+^). (**F**) Kinetic analysis of the cleavage of pre-tRNA^Lys^ by PRORP in presence of TRMT10C-SDR5C1 (4.5 mM Mg^2+^).

Comparably to its effect on ELAC2, TRMT10C-SDR5C1 did not stimulate, but slightly inhibited pre-tRNA 5’-end cleavage by *A. thaliana* PRORP3 (Figure 3B). Furthermore, attempts to replace the TRMT10C-SDR5C1 complex in its mtRNase P function by related methyltransferases, supposed to similarly remold the bound pre-tRNA, failed. Neither *S. cerevisiae* Trm10p nor human TRMT10A, both evolutionarily and structurally related to TRMT10C and able to methylate mitochondrial pre-tRNA^Ile^ *in vitro* (19), rendered pre-tRNA^Ile^ cleavable by human PRORP (Figure 3C). Collectively, all these results suggest a high specificity in the interplay of PRORP with the TRMT10C-SDR5C1 complex, a specificity that appears mainly determined by the direct molecular interactions of PRORP and TRMT10C.

### TRMT10C-SDR5C1 increase the cleavage rate of PRORP

The finding that PRORP alone is able to correctly cleave certain pre-tRNAs, allowed us to quantitatively asses the contribution of TRMT10C-SDR5C1 to the cleavage process by a comparative kinetic analysis. We studied the kinetics of cleavage by the mtRNase P holoenzyme and by PRORP alone under single-turnover conditions (enzyme concentration in excess over substrate). Under these conditions, single rounds of catalysis occur, and product release and substrate-rebinding events do not contribute to the observed reaction rates, thereby putting the focus on the steps after the formation of the initial enzyme-substrate-encounter complexes (which includes the chemical step and possible conformational steps preceding it). We studied the cleavage kinetics of two mitochondrial pre-tRNA model substrates. In the case of the mtRNase P holoenzyme (PRORP + TRMT10C-SDR5C1), we measured a *K*_M(sto)_ of ∼70 nM and a *k*_react_ of ∼1.5 min^-1^ for pre-tRNA^Ala^ (Figure 4A), and a *K*_M(sto)_ of ∼30 nM and a *k*_react_ of ∼1.2 min^-1^ for pre-tRNA^Met^ (Figure 4B). With PRORP alone, we observed an ∼10-to 12-fold lower *k*_react_, but essentially identical *K*_M(sto)_ values for both pre-tRNAs (Figure 4C and 4D), suggesting that the TRMT10C-SDR5C1 complex increases the rate of the conformational rearrangements and/or the chemical step, but not the affinity of PRORP for the substrates. In addition, we examined the mtRNase P-holoenzyme cleavage kinetics of three additional pre-tRNA substrates, pre-tRNA^Ile^, pre-tRNA^Lys^ and pre-tRNA^Val^, for which we observed no cleavage by PRORP alone (Figure 1). Interestingly, *K*_M(sto)_ and *k*_react_ were in the same order of magnitude of those of pre-tRNA^Ala^ and pre-tRNA^Met^, or showed an even higher *k*_react_ in the case of pre-tRNA^Ile^ (Figure 4E and 4F), indicating that the pre-tRNAs that are not cleaved by PRORP alone are not inherently poorer substrates for mtRNase P.

### PRORP binds pre-tRNAs with high affinity

The low nanomolar *K*_M(sto)_ of PRORP, independent of and unchanged by TRMT10C-SDR5C1, suggests that PRORP itself binds pre-tRNAs with high affinity and that the TRMT10C-SDR5C1 complex does not significantly contribute to catalytic efficiency by enhancing PRORP’s substrate affinity. We used bio-layer interferometry to directly study pre-tRNA binding by the subunits of mtRNase P. The method allows the label-free kinetic characterization of macromolecular interactions in real time. One of the binding partners is immobilized on a sensor tip, through which a beam of white light is projected, and the interference pattern of light reflected at the surface and an internal reference layer is recorded. The binding of interacting macromolecules changes the interference pattern, and the shift, expressed in nanometers, is thus a measure of binding. We immobilized the pre-tRNA on the sensor and followed the association of the different proteins and their dissociation upon dilution. We assayed the binding of TRMT10C, SDR5C1, and PRORP, to pre-tRNA^Ala^, pre-tRNA^His^, pre-tRNA^Lys^, and pre-tRNA^Val^, and derived the association (*k*_on_) and dissociation (*k*_off_) rate constants, as well as the dissociation constant (*K*_D_) for each combination (Table 1). TRMT10C and PRORP showed pre-tRNA binding with *K*_D_ values in the low nanomolar range, whilst we observed no binding of SDR5C1 to any of the tested pre-tRNAs, confirming that SDR5C1 alone is not able to bind pre-tRNAs directly, as previously observed (19) and consistent with the structural information (22). The addition of SDR5C1 to TRMT10C nevertheless slightly lowered the observed *K*_D_ for all tested pre-tRNAs, mostly due to a reduction of the *k*_off_, suggesting that the interaction with SDR5C1 somewhat stabilizes the binding of TRMT10C to the pre-tRNA substrate; still, this stabilizing effect is insufficient to explain the dramatic stimulation of TRMT10C’s methylation activity upon supplementation with SDR5C1 (19). Finally, the *K*_D_ of PRORP for pre-tRNAs was in a similar range of that of TRMT10C and of PRORP’s *K*_M(sto)_ for the cleavage reactions, and did not correlate with the ability to cleave or to do not cleave a given pre-tRNA substrate on its own. The similarly high affinity of TRMT10C-SDR5C1 and PRORP for pre-tRNAs unfortunately prevented analogous binding studies with the mtRNase P holoenzyme, and mixtures of all three proteins did not allow any meaningful bio-layer interferometry measurements.

**Table 1.**
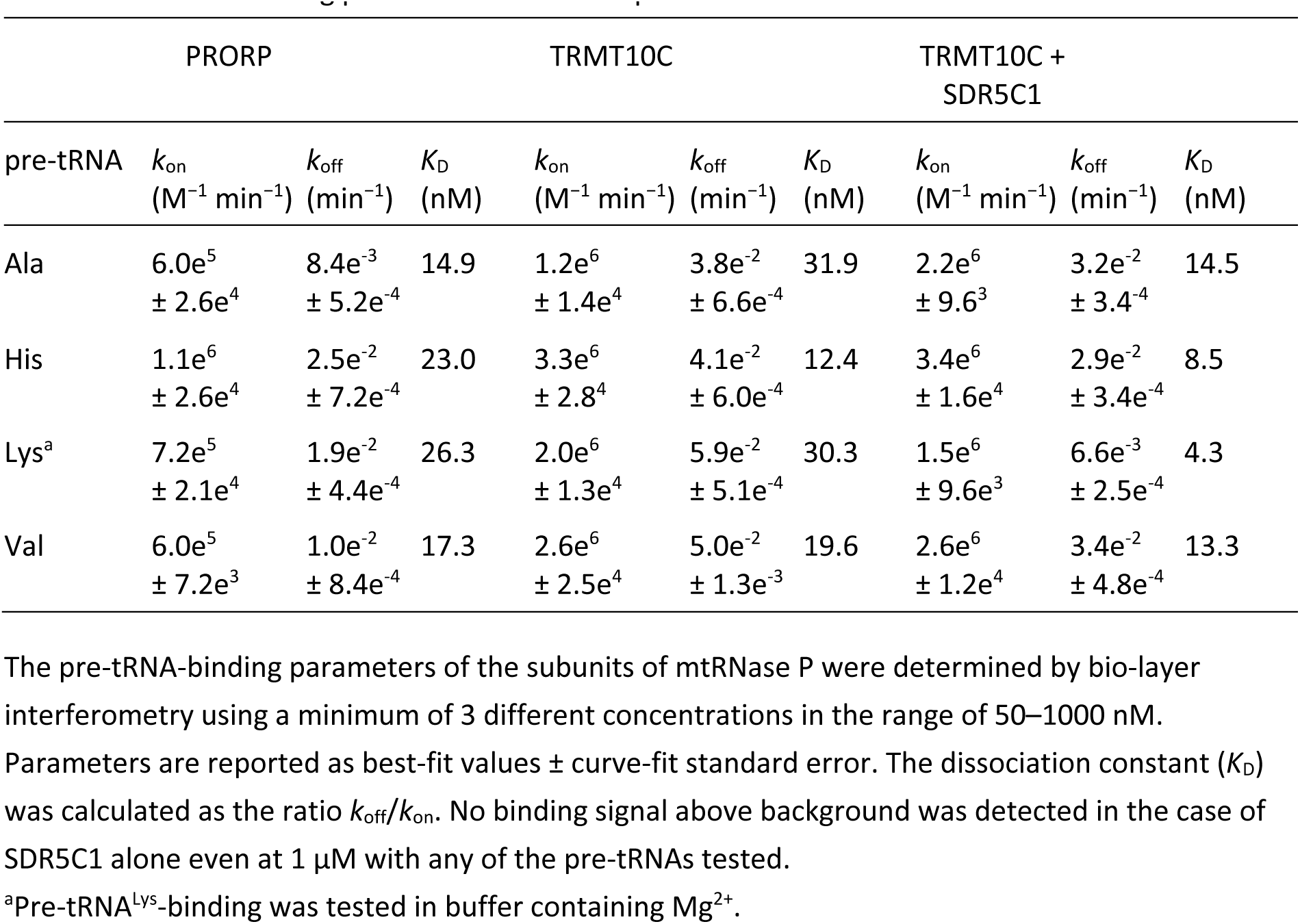
Pre-tRNA binding parameters of the components of mtRNase P.

### PRORP does not require TRMT10C-SDR5C1 or pre-tRNA to properly coordinate Mg^2+^ in its active site

Our comparative kinetic analysis and pre-tRNA-binding studies suggest that, while PROPR is able to efficiently bind pre-tRNAs on its own, its interaction with TRMT10C-SDR5C1 is primarily required for some kind of “activation” through structural changes in – and/or by repositioning of – its nuclease domain leading to significant rate enhancements and making some pre-tRNAs cleavable in the first place. Such scenario appears at first sight consistent with a model that had been proposed on the basis of the crystal structures of PRORP fragments (28,29). In these structures the active site of PRORP appeared distorted, unable to coordinate Mg^2+^, and the authors proposed that the interaction with TRMT10C-SDR5C1 could remold the active site to restore Mg^2+^-coordination by PRORP and hence its endonuclease activity. Our finding that PRORP alone is able to cleave some pre-tRNAs, even though with reduced efficiency, appears difficult to reconcile with a strict requirement for TRMT10C-SDR5C1 to enable the coordination of the catalytic metal ions. Therefore, we used iron-mediated hydroxyl radical cleavage (36) to directly probe whether PRORP alone is able to coordinate divalent metal ions in its active site. When PRORP was incubated with Fe(II) and a reducing agent, the protein underwent partial fragmentation producing polypeptides of about 50 kDa, 40 kDa, 20 kDa and 12 kDa (Figure 5A, lanes 4, 5, 6; the smaller C-terminal fragments could only be visualized by western blotting using an antibody against the C-terminal His-tag). This pattern is consistent with cleavage close to D409 and to one of the other three aspartate residues (D478, D479, D499), all four presumably involved in metal-ion coordination (23,28,29). To confirm the specificity of the iron-mediated cleavage, we used substitution variants D479N and D499N; both aspartates are directly involved in metal-ion coordination in *A. thaliana* PRORP1 (23), but were reported to be rearranged and unavailable for that purpose in human PRORP (28). Both substitutions prevented iron-mediated cleavage (Figure 5A), confirming the involvement of the two aspartates in metal-ion coordination in human PRORP. An identical pattern and extent of iron-mediated fragmentation was observed when equimolar amounts of pre-tRNA^Ile^ or the TRMT10C-SDR5C1 complex, or both, were added to the reaction (Figure 5B, lanes 4, 6 and 7). These results demonstrate that PRORP is able to coordinate metal ions in its active site irrespectively of an interaction with TRMT10C-SDR5C1 or pre-tRNA, consistent with its ability to also cleave at least some pre-tRNAs independently of TRMT10C-SDR5C1.

**Figure 5.**
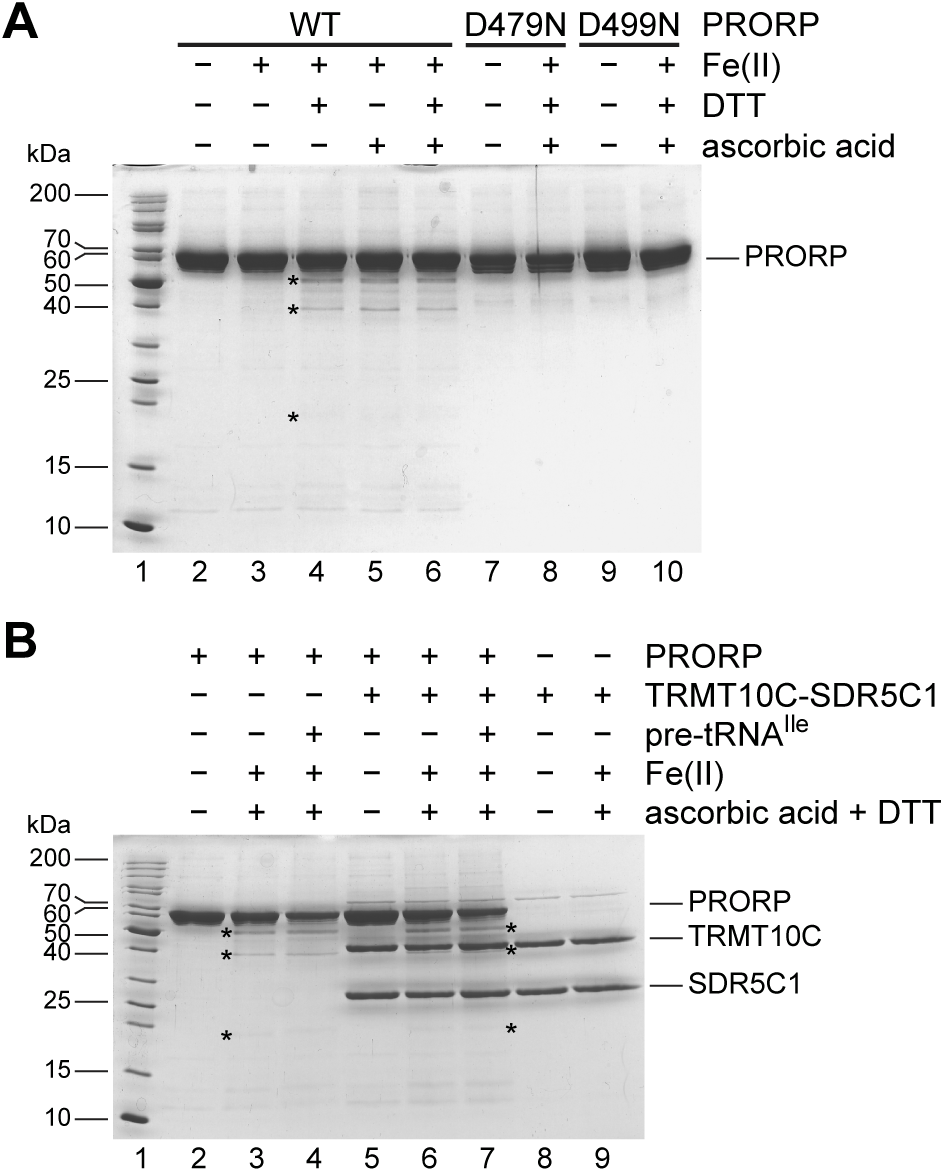
Metal-ion coordination in the active site of PRORP. (**A**) PRORP was subjected to iron-mediated hydroxyl radical cleavage. The wild type protein, or substitution variants D479N and D499N, were incubated with Fe(II), DTT and/or ascorbic acid, and cleavage products were resolved by SDS-PAGE; the gel was stained with Coomassie blue. Specific cleavage products in lanes 4–6 are indicated by asterisks (typically, only a small fraction of the protein is cleaved by this procedure, as previously demonstrated for other, well-established metalloenzymes (36,57)); the barely visible ∼20 kDa fragment and the second, smaller complementary C-terminal fragment (not detectable by direct staining) were confirmed by western blotting. The molecular weight of selected size markers (lane 1) is indicated to the left. The position of full-length PRORP is indicated to the right. (**B**) PRORP was subjected to iron-mediated hydroxyl radical cleavage in the presence of pre-tRNA^Ile^ (lane 4) and/or the TRMT10C-SDR5C1 complex (lanes 7 and 6). Specific cleavage products in lanes 3, 4 and 6, 7, are indicated by asterisks (note that the ∼40-kDa PRORP fragment migrates very close to TRMT10C in lanes 6 and 7, and can only be seen when zooming into the relevant area). The positions of full-length PRORP, TRMT10C and SDR5C1 are indicated to the right.

Higher Mg^2+^ concentrations could be expected to at least partially rescue an impaired coordination of Mg^2+^ in the active site, hence the Mg^2+^ optimum for cleavage by PRORP alone could be higher than that of the holoenzyme if such impairment were the case. Therefore, we also compared the Mg^2+^ optimum for cleavage by PRORP alone with that of the mtRNase P holoenzyme for three pre-tRNA model substrates. For pre-tRNA^Ala^ and pre-tRNA^Glu^ the highest cleavage rate was reached at the same Mg^2+^ concentration with both, PRORP and mtRNase P, albeit the optima for the two pre-tRNAs differed slightly (3 vs. 4.5 mM; Figure 6A and 6B). In the case of pre-tRNA^Met^, the Mg^2+^ optimum of PRORP alone was twofold higher than that of mtRNase P (6 vs. 3 mM; Figure 6C). These results suggest that the affinity of human PRORP for divalent metal ions is not generally and systematically altered by the TRMT10C-SDR5C1 complex, although the Mg^2+^ optimum of PRORP and the holoenzyme may slightly differ in some cases.

**Figure 6.**
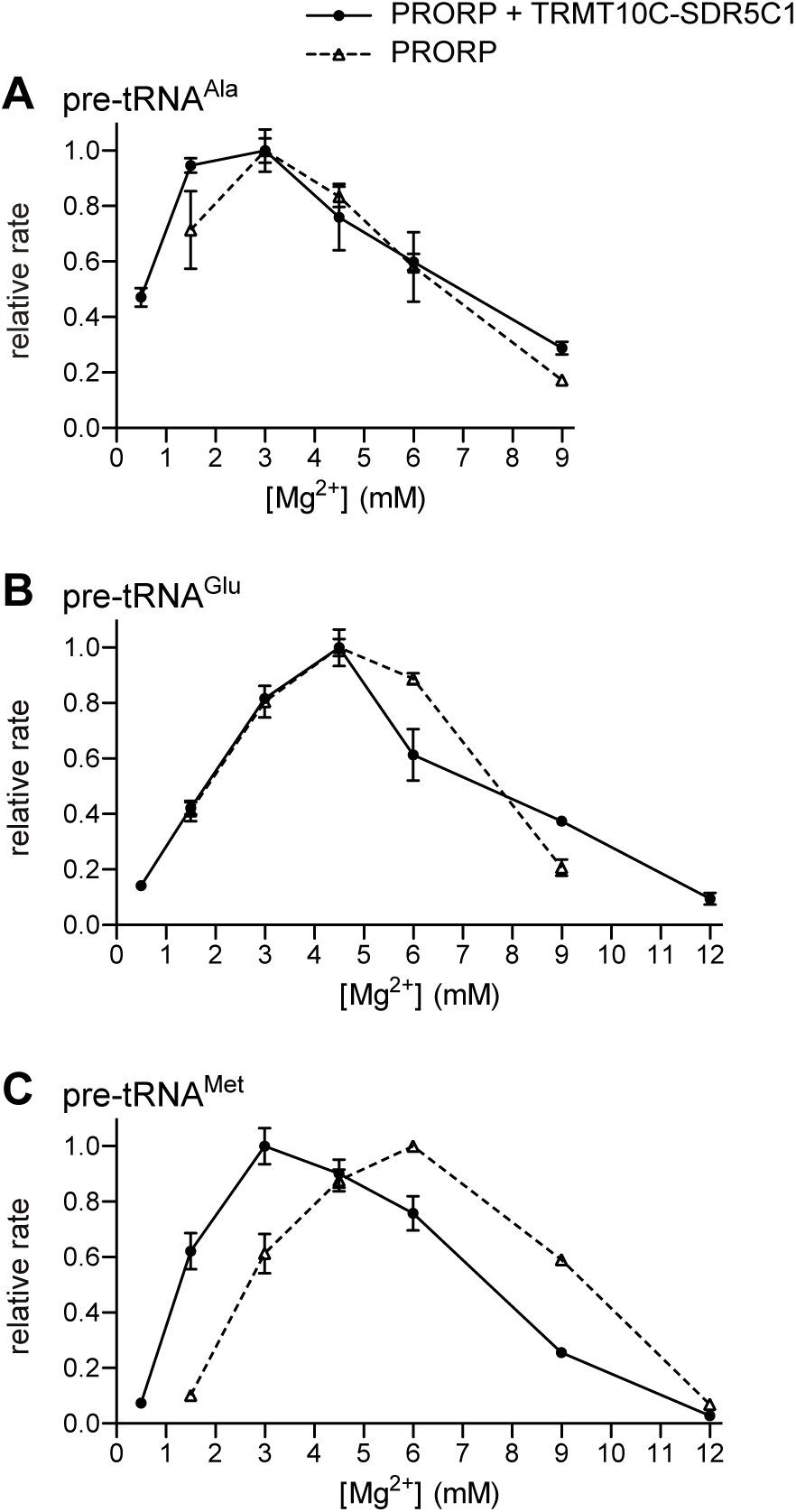
Mg^2+^ optimum of PRORP compared to that of the mtRNase P holoenzyme. Cleavage kinetics were performed with PRORP alone and in the presence of TRMT10C-SDR5C1 (mtRNase P holoenzyme), at different Mg^2+^ concentrations under single-turnover conditions. Pseudo first-order cleavage rates (*k*obs or *k*obs*) were derived and normalized to the highest rate measured. Relative rates represent the mean ± SEM of at least 3 replicates. Cleavage by PRORP alone at the lowest and/or highest concentration shown for the mtRNase P holoenzyme was often too weak to derive a corresponding cleavage rate. (**A**) Mg^2+^ optimum for the cleavage of pre-tRNA^Ala^. (**B**) Mg^2+^ optimum for the cleavage of pre-tRNA^Glu^. (**C**) Mg^2+^ optimum for the cleavage of pre-tRNA^Met^.

### Cleavage chemistry is the rate-limiting step in mtRNase P catalysis

To complete our studies of mtRNase P catalysis, we also performed experiments addressing the early and late steps of catalysis, i.e., steps preceding or following the chemical (cleavage) step, respectively. First, we studied the multiple-turnover kinetics of cleavage by mtRNase P to see, by comparison with single-turnover kinetics, whether product release possibly is a rate-limiting step, like in the case of catalysis by the bacterial and yeast nuclear RNA-based enzymes (47–49), or not, like found for *A. thaliana* PRORP1 (50). We again used saturating concentrations of TRMT10C-SDR5C1 and determined *K*_M_ and *k*_cat_ for two pre-tRNA model substrates also analyzed under single-turnover conditions (see above and Figure 4). For both, pre-tRNA^Met^ and pre-tRNA^Lys^, we measured a *k*_cat_ almost identical to the *k*_react_ obtained under single-turnover conditions and a *K*_M_ only slightly higher than the *K*_M(sto)_ (Figure 7; compare to Figure 4D and 4F). These results essentially rule out that product release is the rate-limiting step under multiple-turnover conditions and suggest that either the chemical step itself or a step before is rate-limiting.

**Figure 7.**
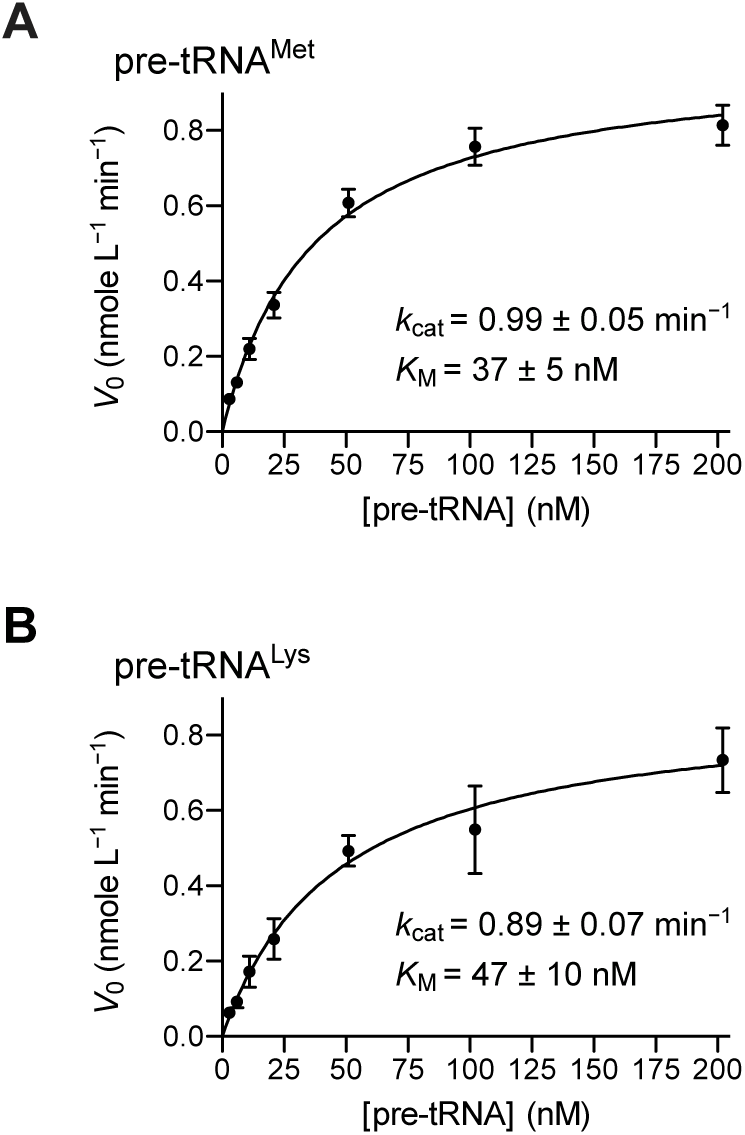
Multiple-turnover kinetics of pre-tRNA cleavage by mtRNase P. Multiple-turnover kinetic analyses of pre-tRNA cleavage were performed with 1 nM PRORP in the presence of an excess of TRMT10C-SDR5C1 complex and with varying concentrations of pre-tRNA (4.5 mM Mg^2+^). Initial velocities of cleavage (*V*0; see Materials and Methods) were plotted against the pre-tRNA concentration. Data points are the mean ± SEM of at least 5 replicates. Derived kinetic constants *k*cat and *K*M are inserted into each graph. (**A**) Multiple-turnover kinetic analysis of the cleavage of pre-tRNA^Met^ by mtRNase P. (**B**) Multiple-turnover kinetic analysis of the cleavage of pre-tRNA^Lys^ by mtRNase P.

Finally, we performed pulse-chase experiments as introduced by the Uhlenbeck lab in their studies of hammerhead ribozymes (34), to shed light on the relationship between the dissociation rate (*k*_−1_) of pre-tRNA and mtRNase P, and the rate constant of the chemical step (*k*_2_). Under single-turnover conditions with an excess of enzyme, cleavage reactions were quenched after a short period of reaction progress, either by addition of an excess of unlabeled substrate or by dilution with reaction buffer. Assuming all substrate is enzyme-bound at the timepoint of the chase, an unaltered progress of the reaction would indicate that *k*_2_ exceeds *k*_−1_ by far, whereas an immediate halt would indicate the reverse, i.e., *k*_−1_ ≫ *k*_2_. MtRNase P showed an intermediate behavior, with some cleavage continuing to occur during the chase (Figure 8). This kinetic behavior indicates that the rate of substrate dissociation and of chemistry are in a similar order, with *k*_2_ probably exceeding *k*_−1_, based on the dissociation rates measured for PRORP alone by bio-layer interferometry (Table 1; substrate dissociation still has to be assumed to be slightly faster under assay conditions due to the higher temperature used; 30 versus 21 °C)).

**Figure 8.**
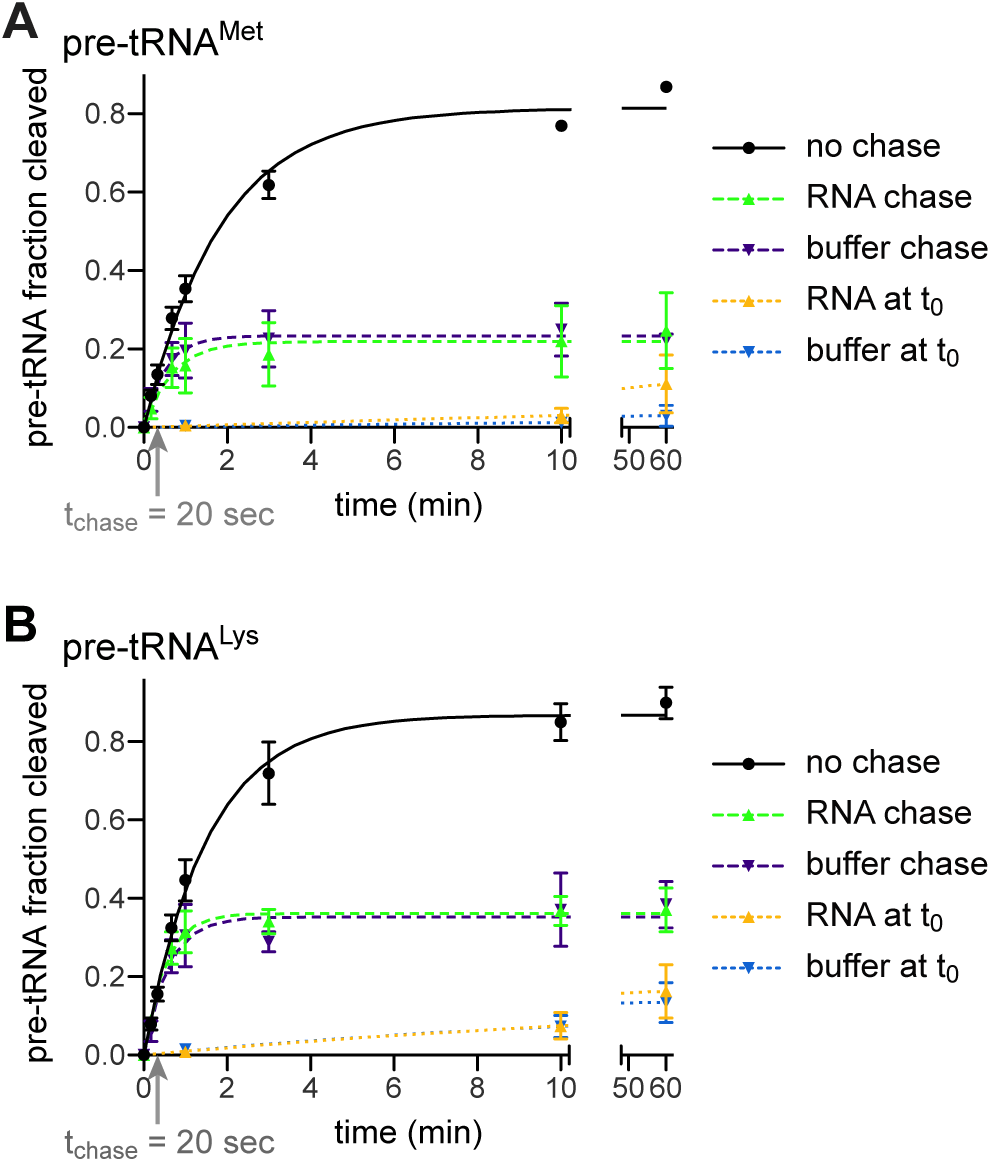
Pulse-chase kinetics of pre-tRNA cleavage by mtRNase P. Trace amounts of pre-tRNA were cleaved by an excess of mtRNase P (PRORP + TRMT10C-SDR5C1) at 4.5 mM Mg^2+^, and the reactions were quenched (“chase”) after 20 seconds, either by addition of a 1000-fold excess of unlabeled substrate, or by 200-fold dilution with reaction buffer. Aliquots were withdrawn and the reaction stopped at the indicated timepoints. Samples were analyzed by gel electrophoresis and phosphorimaging, and the cleaved fraction of pre-tRNA was determined by image analysis. Control reactions without a “chase” and with the “chase” prior to the start of the reaction (t0) were run in parallel. (**A**) Pulse-chase kinetic analysis of the cleavage of pre-tRNA^Met^ by mtRNase P. (**B**) Pulse-chase kinetic analysis of the cleavage of pre-tRNA^Lys^ by mtRNase P.

## DISCUSSION

Since the initial identification of the protein-only RNase P found in human mitochondria (9), the diversity of its subunits and their additional role in other pathways have obscured the holoenzyme’s mechanism of action and puzzled the field (10). The characterization of homologues of the TRM10 and PRORP families, all active as single proteins, only further underlined the exceptional status of those animal mtRNase P subunits within the respective enzyme families. While the recent cryo-EM structure of human mtRNase P elucidated the interactions of the individual subunits with each other and the pre-tRNA substrate (22), it did not shed light on the mechanistic interplay among the subunits, the dynamic aspects of the assembly of the enzyme-substrate complex, and the kinetics of the reaction. Moreover, the association of pathogenic mutations in the genes encoding the three subunits with largely distinct clinical presentations warrants closer attention to these aspects (51–53).

Here, we sought to more precisely define the mechanism of action of the catalytic subunit PRORP, and to shed light on its interplay with the “accessory” subunits TRMT10C and SDR5C1 in achieving efficient phosphodiester hydrolysis. Unexpectedly, we found that PRORP alone is able to efficiently bind pre-tRNAs and to even cleave some of them, although at an at least 10-fold lower rate than in the presence of TRMT10C-SDR5C1. This suggests a model in which in principle the TRMT10C-SDR5C1 complex and PRORP may each independently bind pre-tRNAs, and exert catalysis: TRMT10C-SDR5C1 by methylating the *N*^1^ of purines found at position 9 in 19 of the 22 mitochondrial tRNAs (19), and PRORP by removing the 5’ extension of at least some mitochondrial pre-tRNAs. Still, only the successive or coincident binding of PRORP and TRMT10C-SDR5C1 to a pre-tRNA, and the resulting direct interaction of TRMT10C with PRORP lead to the efficient removal of the 5’ extension from pre-tRNAs, consistent with the strict requirement of all 3 subunits for efficient mtRNase P function *in vivo* (9). PRORP and TRMT10C-SDR5C1 bind to all tested pre-tRNAs with similar affinity and there is no evidence from the kinetic analyses that either the PRORP·pre-tRNA or the TRMT10C-SDR5C1·pre-tRNA complex represents the preferred pathway to the cleavage-efficient holoenzyme·pre-tRNA complex; the order of events *in vivo* may thus simply depend on the availability and relative stoichiometries of the two players PRORP and TRMT10C-SDR5C1.

In the light of evolution, the vestigial nuclease activity of human PRORP appears not too surprising, given its apparent descent from single-subunit PRORPs (10,18). Remarkably, human PRORP still appears to generally bind pre-tRNAs with similar affinity, regardless of whether it is able to cleave them on its own or not. TRMT10C-SDR5C1 exerts its effect on the cleavage kinetics of PRORP by a rate enhancement rather than by decreasing the *K*_M(sto)_, at least in cases where PRORP alone is also able to cleave the substrate. This suggests that direct interactions with TRMT10C induce rearrangements in and/or a re-positioning of PRORP’s nuclease domain that allow the (more efficient) hydrolysis of the target phosphodiester bond. The interactions of PRORP’s PPR domain with the tRNA “elbow” are probably not affected by the interaction with TRMT10C and, considering the latter’s kinetic effect, they likely remain similar or even identical in the holoenzyme.

Conformational changes as a mechanism in the activation of human PRORP by TRMT10C-SDR5C1 were previously proposed based on the crystal structures of PRORP fragments (28,29) and also discussed in comparison with the more recent cryo-EM structure of the holoenzyme (22). The authors suggested that the active site of PRORP naturally is in an auto-inhibitory conformation, in which crucial aspartates are rearranged or sequestered, and thereby unavailable for Mg^2+^ coordination. The interaction with TRMT10C-SDR5C1 was suggested to remold this quasi-physiologically disordered catalytic domain to enable the coordination of Mg^2+^ in PRORP’s active site. However, our experiments show that PRORP can actually coordinate divalent metal ions irrespectively of the presence of TRMT10C-SDR5C1 or pre-tRNA, and the ability of PRORP to cleave a subset of pre-tRNAs on its own also demonstrates that it properly coordinates the essential cofactor Mg^2+^ in its active site; in addition, neither we, nor others (54) observed a systematic enhancement of the Mg^2+^ affinity of PRORP by TRMT10C-SDR5C1. This suggests that the rearrangements observed in the crystals of PRORP are either an artifact of the used fragments (28,29) and do not represent a physiologically relevant conformation of the nuclease, or they are a consequence of the absence of metal ions in the crystals. The latter appears consistent with the absence of metal ion 1 in the cryo-EM structure and a D478 side chain that is tilted away (“rearranged”) from its metal coordinating position (22) when compared to the corresponding D474 side chain in the *A. thaliana* PRORP1 crystal structure, which contains both metal ions (23). Still, the disparate steric arrangement of D478 and D479 in the two PRORP crystals also casts doubts on the significance of the structures, and particularly the rearrangements of the central domain could be the artifactual result of the truncated PPR domain of the fragments (28,29); notably, the PRORP fragments used for crystallization were catalytically inactive even in the presence of TRMT10C-SDR5C1. In the end, our results demonstrate that the nuclease domain of PRORP is in principle functional (on its own) and a restoration of the ability to coordinate Mg^2+^ is apparently not the principle underlying the “activating” effect of TRMT10C-SDR5C1.

The favorable positioning or locking-into-position of PRORP’s nuclease domain on the tRNA’s acceptor stem, and its proper adjustment to the cleavage site rather appear to be the major conformational adjustment(s) that are induced by the interaction with TRMT10C; simultaneously, TRMT10C interacts with PRORP’s PPR domain and holds it in place (22). Although the ancient binding mode of single-subunit PRORPs has apparently been partially retained by the human homolog, as indicated by the reasonably high affinity with which it binds pre-tRNAs, it is no longer catalytically productive, depending on the respective substrate, leading to no, low, or aberrant cleavage activity by PRORP alone. Substantial miscleavage of *T. thermophilus* pre-tRNA^Gly^ by PRORP alone, but not by the mtRNase P holoenzyme, suggests that the acceptor stem, the cleavage site and/or the 5’-leader are positioned differently in the active site of PRORP’s metallonuclease domain, depending on whether it interacts with the pre-tRNA only or a TRMT10C-SDR5C1·pre-tRNA complex. The subtle rearrangement of the TRMT10C-SDR5C1-bound tRNA structure observed in the cryo-EM structure (22), might be irrelevant for the cleavage process in this scenario, and the direct interactions of TRMT10C with PRORP’s nuclease domain appear to be the major driver of the conformational adjustments that boost the catalytic activity of PRORP.

Pulse-chase experiments suggest that some step before cleavage limits reaction velocity, which appears consistent with the suggested conformational adjustments of the nuclease domain on the substrate through its interaction with TRMT10C. Comparison of the kinetics of cleavage by the mtRNase P holoenzyme (PRORP in the presence of an excess of TRMT10C-SDR5C1) finally revealed that the maximal rate constant (*k*_react_) determined under single-turnover conditions and the turnover number (*k*_cat_) derived from multiple-turnover analyses were almost identical for the two substrates analyzed. Thus, unlike in the cleavage by bacterial or yeast nuclear RNase P (RNA-based enzymes) (47–49), but similar to that of *A. thaliana* PRORP1 (50), product release is not a rate-limiting step in the catalysis by human mtRNase P. However, the mechanistic basis for the lack of product inhibition remains unclear.

Although we did not directly address the methylation or cleavage kinetics of TRMT10C-SDR5C1 in this study, preliminary attempts to detect a multiple-turnover ability failed, consistent with observations previously reported by others (46). However, such a lack of multiple turnover capacity is surprising, given that the dissociation rate constants for TRMT10C-SDR5C1 were on average not significantly lower than those for PRORP under the same conditions. Thus, the mechanistic basis of the lacking multiple-turnover ability and how the release of products (5’-processed and/or methylated pre-tRNAs) from TRMT10C-SDR5C1 is achieved, remain unclear.

In an independent mechanistic study of human mtRNase P based on kinetic and binding experiments and available as a preprint, the binding affinity of the mtRNase P holoenzyme for pre-tRNA, as measured by fluorescence anisotropy, was reported to be greater than that of the separate subunits (54). Based on that finding and the kinetic enhancement of cleavage by increasing concentrations of TRMT10C-SDR5C1, the authors proposed that TRMT10C-SDR5C1 contributes to cleavage by increasing the binding affinity of PRORP in addition to a rate enhancement. Although we could not directly quantify pre-tRNA binding of the holoenzyme, our kinetic data do not indicate such an effect, as the *K*_M(sto)_ of PRORP in the cleavage of two different pre-tRNA substrates was unaffected by TRMT10C-SDR5C1. However, the pre-tRNA affinity of PRORP that we measured by bio-layer interferometry was substantially (almost 100-fold) higher than in their case and they also did not observe cleavage by PRORP alone. While the latter difference might be attributable to the specific bacterial pre-tRNA substrate they used, the affinity of PRORP for pre-tRNAs cleaved by PRORP alone and for those requiring TRMT10C-SDR5C1 was similar in our hands.

The TRMT10C-SDR5C1 complex was previously suggested to also stimulate pre-tRNA cleavage by mitochondrial RNase Z (ELAC2) (46). However, we could not reproduce this observation, but rather found a consistent inhibition of the RNase Z activity of ELAC2 under all the conditions tested. In addition, ELAC2 is per se kinetically efficient and comparable to the mtRNase P holoenzyme rather than PRORP alone, further questioning a physiological need for an accessory subunit like TRMT10C-SDR5C1. Finally, in contrast to the mtRNase P subunits, ELAC2 is dually localized and responsible for the endonucleolytic 3’-end processing of nucleus-encoded tRNAs too (5,6,55), evidently in the absence of TRMT10C-SDR5C1.

As discussed above, TRMT10C-SDR5C1 appears to enable or stimulate mtRNase P cleavage activity by adjusting PRORP’s nuclease domain and active site to the commonly bound pre-tRNA via specific protein-protein interactions. The pre-tRNAs that are cleaved by PRORP alone might have a more favorable orientation of the domains due to their stem and loop lengths, or nucleotides at key positions that by themselves favor the correct positioning in the active site. However, due to the limited number of mitochondrial pre-tRNA substrates, it is currently impossible to derive a consensus for the features that make a pre-tRNA cleavable by PRORP alone, though at low efficiency. Alternatively, the tRNAs that are processed by PRORP alone may benefit from a particular flexibility, which allows them to transiently assume conformations more favorable for cleavage. Indeed, human mitochondrial pre-tRNAs have rather unstable structures due to an elevated number of A-U base pairs, non-canonical base pairs and mismatches in their stems, and the frequent lack of the conserved tertiary interactions, and, as a consequence, are predicted to assume multiple alternative conformations in solution (Vilardo, Wolfinger, Hofacker and Rossmanith, unpublished analysis). When we used *T. thermophilus* pre-tRNA^Gly^, a fully canonical tRNA characterized by G-C rich stems and therefore a stable and probably more rigid structure, this substrate only partially adopted the canonical binding mode, as inferred from 50% miscleavage. Thus, TRMT10C-SDR5C1 acts as an adaptor between pre-tRNA and PRORP, locking the substrate in the conformation leading to correct cleavage by the endonuclease subunit. The mitochondrial methyltransferase TRMT10C-SDR5C1, which is only found in metazoans (56), has apparently been recruited as a general accessory factor to safeguard proper cleavage of the structurally degenerating pre-tRNA substrates by PRORP, like an extra recognition-handle beyond the direct contacts of PRORP with the pre-tRNA. The cooperation might have resulted in the catalytic degeneration of PRORP and developed into a specific catalytic dependency on the interaction partner.

## ACKNOWLEDGEMENTS

We acknowledge the contribution of Nadia Brillante of a few pre-tRNA processing experiments, and the technical support of Fatinah El-Isa and Tobias Pober with some of the enzyme kinetic experiments.

## FIGURES

## TABLES

